# Optical manipulations reveal strong reciprocal inhibition but limited recurrent excitation within olfactory bulb glomeruli

**DOI:** 10.1101/2021.07.21.453150

**Authors:** Joseph D. Zak, Nathan E. Schoppa

**Affiliations:** Department of Physiology & Biophysics, University of Colorado Anschutz Medical Campus, Aurora, CO 80045; Neuroscience Program, University of Colorado Anschutz Medical Campus, Aurora, CO 80045; Department of Molecular & Cellular Biology, Harvard University, Cambridge, MA 02138

## Abstract

The local circuitry within olfactory bulb glomeruli filters, transforms, and facilitates information transfer from olfactory sensory neurons to bulb output neurons. Two key elements of this circuit are glutamatergic tufted cells (TCs) and GABAergic periglomerular (PG) cells, both of which actively shape mitral cell activity and bulb output. A subtype of TCs, the external tufted cells (eTCs), can synaptically excite PG cells, but there are unresolved questions about other aspects of the glomerular connections, including the extent of connectivity between eTCs and the precise nature of reciprocal interactions between eTCs and PG cells. We combined patch-clamp recordings in OB slices and optophysiological tools to investigate local functional connections within glomeruli. When TCs were optically suppressed, we found a large decrease in excitatory post-synaptic currents (EPSCs) in “uniglomerular” PG cells that extend dendrites to one glomerulus, indicating that TC activation was required for most excitation of these PG cells. However, TC suppression had no effect on EPSCs in eTCs, arguing that TCs make few, if any, direct excitatory synaptic connections onto eTCs. The absence of synaptic connections between eTCs was also supported by recordings in eTC pairs. Lastly, we show using similar optical suppression methods that PG cells that express GAD65, mainly uniglomerular PG cells, provide strong inhibition onto eTCs. Our results indicate that the local network of TCs form potent reciprocal synaptic connections with GAD65-expressing uniglomerular PG cells but not other TCs. This configuration favors local inhibition over recurrent excitation within a glomerulus, limiting information transfer to downstream cortical regions.

## Introduction

Olfactory bulb (OB) glomeruli not only serve as a relay between the sensory periphery and the olfactory cortex but are also the site of initial information processing in the olfactory system. Glomeruli are innervated by multiple cell types that facilitate the detection and discrimination of odors through interconnected networks of excitatory and inhibitory neurons (Nagayama et al., 2014; Takahashi et al., 2016; Burton, 2017). A class of excitatory interneurons known as external tufted cells (eTCs) are a central element that controls both excitation and inhibition within glomeruli. For example, eTCs can directly excite a subset of GABAergic periglomerular (PG) cells through glutamatergic synaptic contacts (Hayar et al., 2004a). These PG cells, which have functionally been defined as Type II cells (Shao et al., 2009), account for ∼70% of the total number of PG cells. At the same time, eTCs also excite OB output mitral cells (MCs) (De Saint Jan et al., 2009; Najac et al., 2011; Gire et al., 2012). This eTC-to-MC excitation, which appears to result from glutamate “spillover” at eTC-to-PG cell dendrodendritic synapses (Gire et al., 2019), accounts for the majority of the excitatory charge in MCs following stimulation of olfactory sensory neurons (OSNs; Gire et al., 2012; Vaaga and Westbrook, 2016). Between glomeruli, laterally projecting short axon cells bidirectionally modulate activity at neighboring glomeruli through neurotransmitter co-release and electrical coupling (Aungst et al., 2003; Maher and Westbrook, 2008; Banerjee et al., 2015).

In spite of progress in understanding the functional connectivity of glomerular neurons, there remain numerous unanswered questions. For example, while eTCs can clearly excite Type II PG cells, their connectivity onto other eTCs is not well-resolved. Because of the high sensitivity of eTCs to OSN input (De Saint Jan et al., 2009; Gire et al., 2012), eTC-to-eTC excitation could mediate an important type of recurrent excitation within a glomerulus following OSN stimulation that could impact output neuron activity. Ultrastructural studies have provided somewhat differing perspectives on whether there are dendrodendritic excitatory synapses within glomeruli (Kosaka and Kosaka, 2005; Bourne and Schoppa, 2017), potential eTC-to-eTC contacts, and whether eTCs form synapses with each other has not been well-examined using physiological methods. The role of eTCs in exciting PG cells is also not completely understood, especially as it relates to how much eTCs contribute to excitation of Type II PG cells versus other sources. For example, PG cells can be excited directly by MCs, (Najac et al., 2015) as well as centrifugal inputs from olfactory cortical regions (Boyd et al., 2012; Markopoulos et al., 2012). A last point pertains to the connections from PG cells back onto eTCs. Prior studies have established that there are different subtypes of morphologically distinct PG cells that can be defined by the isoform of glutamate decarboxylase (GAD) that they express (Parrish-Aungst et al., 2007; Kiyokage et al., 2010), either GAD65 or GAD67. It is not clear which of these subtypes provide most of the inhibition onto eTCs.

We studied the local functional connectivity within glomeruli, first, using an optogenetic silencing approach in cholecystokinin (CCK)-Cre mice. CCK is highly expressed in OB tufted cells (TCs) of different subtypes, including eTCs (Sun et al., 2020), and thus these mice provided a convenient tool to examine the functional connectivity from TCs onto different cell types. Second, we used dual whole-cell recordings that included an eTC and either another eTC or a PG cell. These experiments provided evidence that both TCs in general, as well as eTCs in particular, in fact do not form direct excitatory synaptic connections onto other eTCs, even though they potently excite PG cells. Also, TC activation underlies most excitation of Type II PG cells. As a final element of our study, we used recordings in GAD65-Cre mice to provide evidence that GAD65-expressing PG cells mediate a major source of inhibition onto eTCs. Together, our findings support a model in which the local network of TCs form potent reciprocal connections with GAD65-expressing GABAergic interneurons but not other TCs.

## Results

### Optical suppression of CCK-expressing cells

In the OB, the peptide hormone cholecystokinin (CCK) is selectively expressed in TCs (including eTCs), but not MCs (Seroogy et al., 1985; Economo et al., 2016; Sun et al., 2020). We first confirmed CCK expression and spatial distributions by generating CCK-tdTomato mice (see Methods) (Chhatwal et al., 2007; Madisen et al., 2012). In OB slices from reporter mice, tdTomato expression was largely confined to the inner half of the glomerular layer and outer portion of the external plexiform layer (**Figure 1A**), consistent with prior descriptions of the spatial distribution of CCK expression (Sun et al., 2020). These TCs included eTCs, which reside in the glomerular layer and lack lateral dendrites, and superficial TCs that have lateral dendrites that extend just below the glomerular layer through the external plexiform layer. We also observed sparse labeling of cells deeper in the bulb, which are likely middle and deep TCs. We then took advantage of this selective expression by crossing CCK-Cre mice with a conditional line for the light-activated chloride pump halorhodopsin (NpHR3.0) to generate CCK-NpHR3.0 animals. These mice allowed us to test how optical suppression of TCs influences excitatory signaling within OB glomeruli. Because we could not differentiate between subtypes of TCs based on CCK expression alone, we refer to the impacted cells upon optogenetic suppression collectively as TCs.

**Figure 1.**
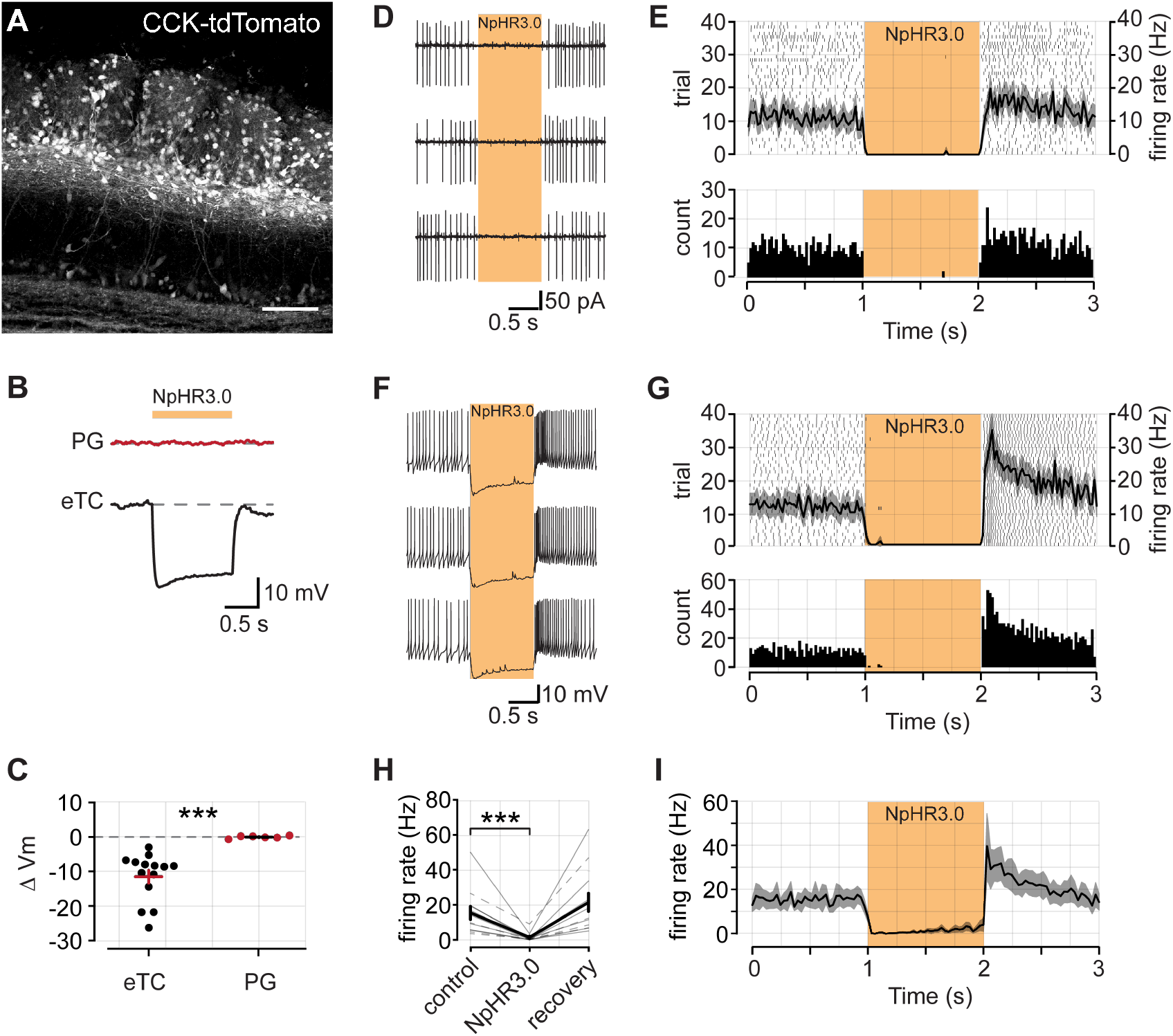
CCK-Cre mice provide a tool to suppress activity specifically in bulbar tufted cells (TCs). **A**. CCK expression in a tdTomato reporter mouse line. CCK is densely expressed in cells in the inner portion of the glomerular layer and outer portion of the external plexiform layer (EPL). Scale bar = 100 μm. **B**. Example membrane voltage responses from a PG cell (top) and an eTC (bottom) in a CCK-NpHR3.0 mouse in response to a 1 s pulse of 590 nm LED light. **C**. Summary data of NpHR3.0 induced membrane voltage changes in eTCs (*n* = 14) and PG cells (*n* = 6). **D**. Example spike recordings from an eTC in LCA configuration. Yellow bar denotes 590 nm light pulse. **E**. *Top*, Raster plot of 40 trials recorded from the same cell as in part *D*. The mean PSTH of all trials (black curve) is superimposed. Shaded area denotes standard error, bin width = 30 ms. *Bottom*, a histogram of all trials. **F-G**. Example spike recordings from an eTC in current-clamp configuration. Plots are the same as *D-E*. **H**. Summary data from combined LCA (*n* = 4; dashed lines) and current-clamp (*n* = 8; solid lines) experiments. Firing rate calculated as the mean in each epoch. Error bars are s.e.m. **I**. Mean PSTH across all recordings (*n* = 12).

To test the efficacy of NpHR3.0 expression, we recorded from visually identified eTCs or PG cells in the glomerular layer of the OB (see Methods). In current-clamp configuration, after a one-second baseline period, we delivered a one-second pulse of 590 nm light to the slice to activate NpHR3.0 (**Figure 1B,D,F**). eTCs were strongly hyperpolarized by illumination (−11.39 ± 1.87 mV, *n* = 14), while there was no effect on PG cell membrane potential (0.10 ± 0.15 mV change, *n* = 6, *P* < 0.001 comparison to eTCs, *Wilcoxon rank sum test*; **Figure 1B-C**). MCs also did not display significant light-evoked hyperpolarizations (−1.24 ± 0.64 mV, *n = 3*). Thus, NpHR3.0 expression appears to be high in TCs but not other OB cells. In terms of preventing action potential firing in eTCs, the light pulses were highly efficacious. Across 12 eTC recordings, conducted in either whole-cell current-clamp (*n* = 8) or loose cell-attached patch configurations (*n* = 4), light nearly eliminated spontaneous spiking (15.47 ± 3.80 Hz baseline epoch vs. 1.39 ± 0.75 Hz LED epoch, *P* = 0.0005, *Wilcoxon signed rank test*; **Figure 1H-I**). In some cells, and shown in the summary data in **Figure 1I**, a small recovery in spiking was observed over the course of the light pulse. This recovery likely stems from a hyperpolarization-activated current that is a hallmark of eTCs (Liu and Shipley, 2008; De Saint Jan et al., 2009), and can be observed in the recordings in **Figure 1B,F**.

### Under baseline conditions, TCs provide most of the direct excitatory input onto Type II PG cells while providing no direct input onto eTCs

With the ability to strongly and selectively suppress TC output, we next examined the contribution of TCs to the excitation of different neurons in the glomerular layer. In this analysis, we focused on both uniglomerular PG cells that extend their dendrites to one glomerulus (Kiyokage et al., 2010), as well as eTCs. Among PG cells, there is a subpopulation of functionally-defined Type II cells that receive direct glutamatergic synaptic input from eTCs (Hayar et al., 2004a; Shao et al., 2009), but the contribution of other glutamatergic cells to their excitation is not well-resolved. At least some PG cells can be directly excited by MCs (Najac et al., 2015) and/or centrifugal inputs (Boyd et al., 2012; Markopoulos et al., 2012). Type II PG cells were identified in our recordings based on the presence of a prolonged barrage of EPSCs in response to electrical stimulation of OSNs (**Figure 2A**).

**Figure 2.**
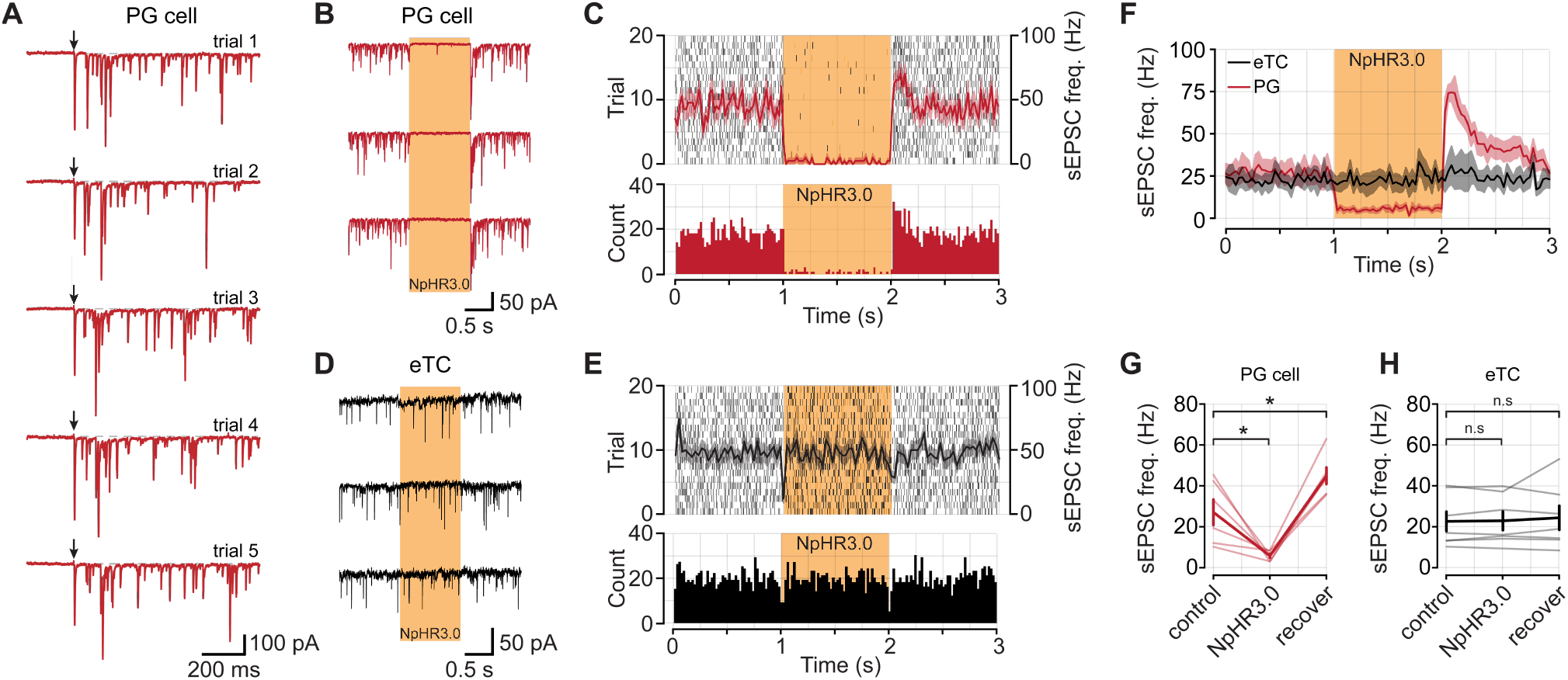
Optical suppression of TCs in CCK-NpHR3.0 mice reduces the frequency of excitatory synaptic inputs onto PG cells but not eTCs. **A**. Subtype of PG cell (Type II versus Type I) for this analysis was determined by applying electrical stimulus pulses to OSN axons (downward arrows). Note the barrage of evoked EPSCs in the voltage-clamped cell (*V*_*hold*_ = –77 mV), characteristic of Type II PG cells. **B**. Example recordings of spontaneous EPSCs (sEPSCs; no stimulation) in a Type II PG cell across three consecutive sweeps. The yellow bar denotes a 590 nm light pulse for 1 s to suppress TC output. **C**. *Top*, Raster plot of 20 trials recorded from the same cell as in *A*. The mean PSTH of all trials is superimposed in red, the shaded area denotes s.e.m. *Bottom*, a histogram of all trials from the raster plot above. **D**. Example sEPSC recordings from an eTC. **E**. Plots are same as *B*, but for the eTC shown in part *D*. **F**. Mean sEPSC PSTH across all PG cells (*n* = 6) and eTCs (*n* = 7). **G**. Summary data from all PG cells. **H**. Summary data from all eTCs.

To examine the contribution of TCs to neural excitation, we recorded fast, AMPA receptor-mediated EPSCs in voltage-clamped Type II PG cells or eTCs (*V_hold_* = -77 mV) in CCK-NpHR3.0 mice and asked what fraction of these events were eliminated by light-evoked silencing of TCs. In the initial analysis, we evaluated spontaneous EPSCs (sEPSCs) that were recorded in the absence of stimulation. We found that optical suppression of TCs resulted in a large, reversible reduction in the frequency of sEPSCs in PG cells (69.65 ± 9.41% reduction, *n* = 6; *P* = 0.031 **Figure 2B,C,F,G**). In contrast, sEPSCs in eTCs were unaffected (2.22 ± 3.81% increase in frequency, *n* = 7; *P* = 0.689; **Figure 2D,E,H**). These data support the idea that TCs are the primary source of input to Type II PG cells under baseline conditions but provide little, if any, direct input to eTCs. Interestingly, in the PG cell recordings, the sEPSC rate during the recovery epoch after light application was somewhat higher than the control, pre-stimulus period (27.07 ± 6.25 Hz baseline vs. 44.89 ± 4.05 recovery; *P* = 0.031; *Wilcoxon signed rank test*; **Figure 2F,G**). This likely reflected eTC spike bursts that were activated by NpHR3.0-induced hyperpolarization-activated currents (Liu and Shipley, 2008; De Saint Jan et al., 2009).

We also measured the amplitudes of the sEPSCs for both PG cells and eTCs (**Figure 3A-C**). During light-evoked suppression of TCs, the amplitudes of the sEPSCs in PG cells were drastically reduced (61.84 ± 9.87% reduction, *P* = 0.031; *Wilcoxon signed rank test;* **Figure 3D**). In these recordings, some smaller amplitude sEPSCs persisted (**Figure 3A,C,D**), which may reflect miniature EPSCs (mEPSCs). mEPSCs are thought to arise from action-potential independent release at a single synapse and therefore should be insensitive to light-driven membrane hyperpolarizations in TCs. In contrast to the potent effects of TC inactivation on sEPSC amplitude in PG cells, TC inactivation had no effect on the sEPSC amplitudes in eTCs (5.18 ± 4.22% reduction, *P* = 0.688; *Wilcoxon signed rank test;* **Figure 3B,C,D**). Finally, to demonstrate that NpHR3.0 activation had no effect on release from OSN axon terminals, we measured evoked monosynaptic EPSCs from OSNs to eTCs (**Figure 3E**). We found no difference in the peak evoked EPSC amplitudes for the same stimulus intensity under control conditions and during slice illumination (372.31 ± 91.51 pA control trials vs. 381.39 ± 88.64 pA NpHR3.0 trials, *n* = 5; *P* = 0.945; *Wilcoxon signed rank test*; **Figure 3F**). Overall, our analysis of EPSC amplitudes was consistent with the conclusions from analyzing sEPSC frequency, that TCs are the major source of excitatory input onto Type II PG cells under baseline conditions but they provide little excitatory input onto eTCs.

**Figure 3.**
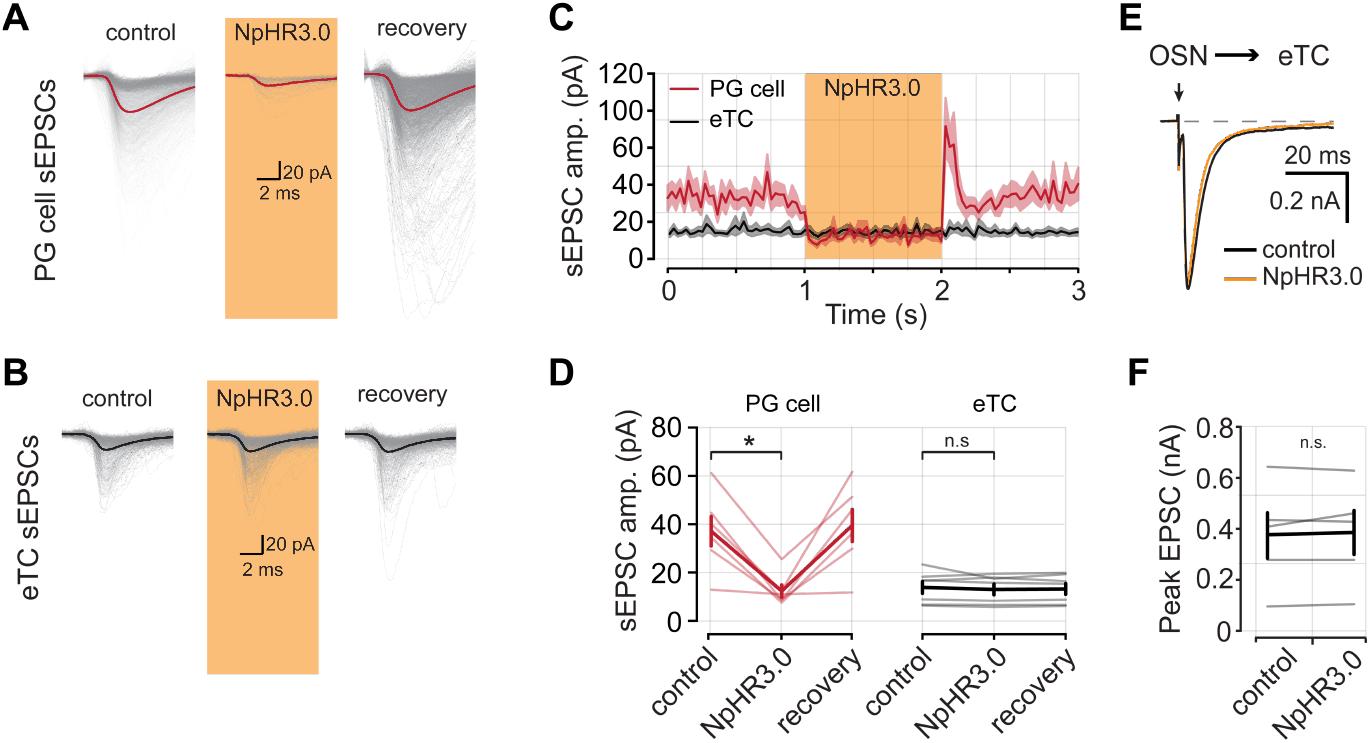
Optical suppression of TCs in CCK-NpHR3.0 mice eliminates large-amplitude sEPSCs in PG cells. **A**. Aligned sEPSCs captured from an example PG cell in each epoch. **B**. Aligned sEPSCs captured from an example eTC. **C**. PSTH of mean sEPSC amplitude across all PG cells and eTCs (*n* = 6 PG cells, *n* = 7 eTCs). **D**. Summary data from all cells in each recording epoch eTCs (61.84 ± 9.87 % reduction PG cells, *P* = 0.031; 5.18 ± 4.22 % reduction eTCs cells, *P* = 0.688; *Wilcoxon signed rank test)*. **E**. Example EPSCs evoked by OSN stimulation recorded from an eTC and during TC inactivation with NpHR3.0. Downward arrow denotes stimulus to OSNs. **F**. Summary of evoked EPSC amplitudes from five eTCs (372.31 ± 91.51 pA control trials vs. 381.39 ± 88.64 pA NpHR3.0 trials; *P* = 0.945; *Wilcoxon signed rank test*).

### Excitation of PG cells driven by OSN stimulation requires activation of TCs

In addition to baseline conditions, we also wanted to know the contribution of different mechanisms of exciting PG cells when the OB network is excited by OSN stimulation. The role of eTCs in particular in exciting PG cells may be unusually high under baseline conditions due to the high level of spontaneous spiking in eTCs (Hayar et al., 2004b). In these studies, we recorded from Type II PG cells and delivered electrical stimulation to OSNs (100 µsec; variable intensity) that was sufficient to generate EPSC barrages on every trial (**Figure 4A**). We then suppressed TC activity using light-evoked activation of NpHR3.0 on alternating blocks of 25 trials (**Figure 4A-C**); in trials with light pulses, OSN stimulation was applied 20 ms after the start of the light pulses. We found that light caused a large ∼75% reduction in the number of evoked EPSCs in PG cells (36.21 ± 3.47 Hz control trials vs. 8.46 ± 1.52 Hz NpHR3.0 trials; *n* = 6; *P* = 0.031; *Wilcoxon signed rank test*; **Figure 4E**). The decrease in evoked EPSCs in PG cells was not the result of changes in the underlying baseline sEPSC rate on control versus NpHR3.0 trials, since these baseline rates were not different (12.93 ± 4.57 Hz control trials vs 17.31 ± 5.08 Hz NpHR3.0 trials; *P* = 0.094; *Wilcoxon signed rank test;* **Figure 4F**). Thus, our results provide good evidence that activation of CCK-expressing TCs was required for most synaptic excitation of Type II PG cells.

**Figure 4.**
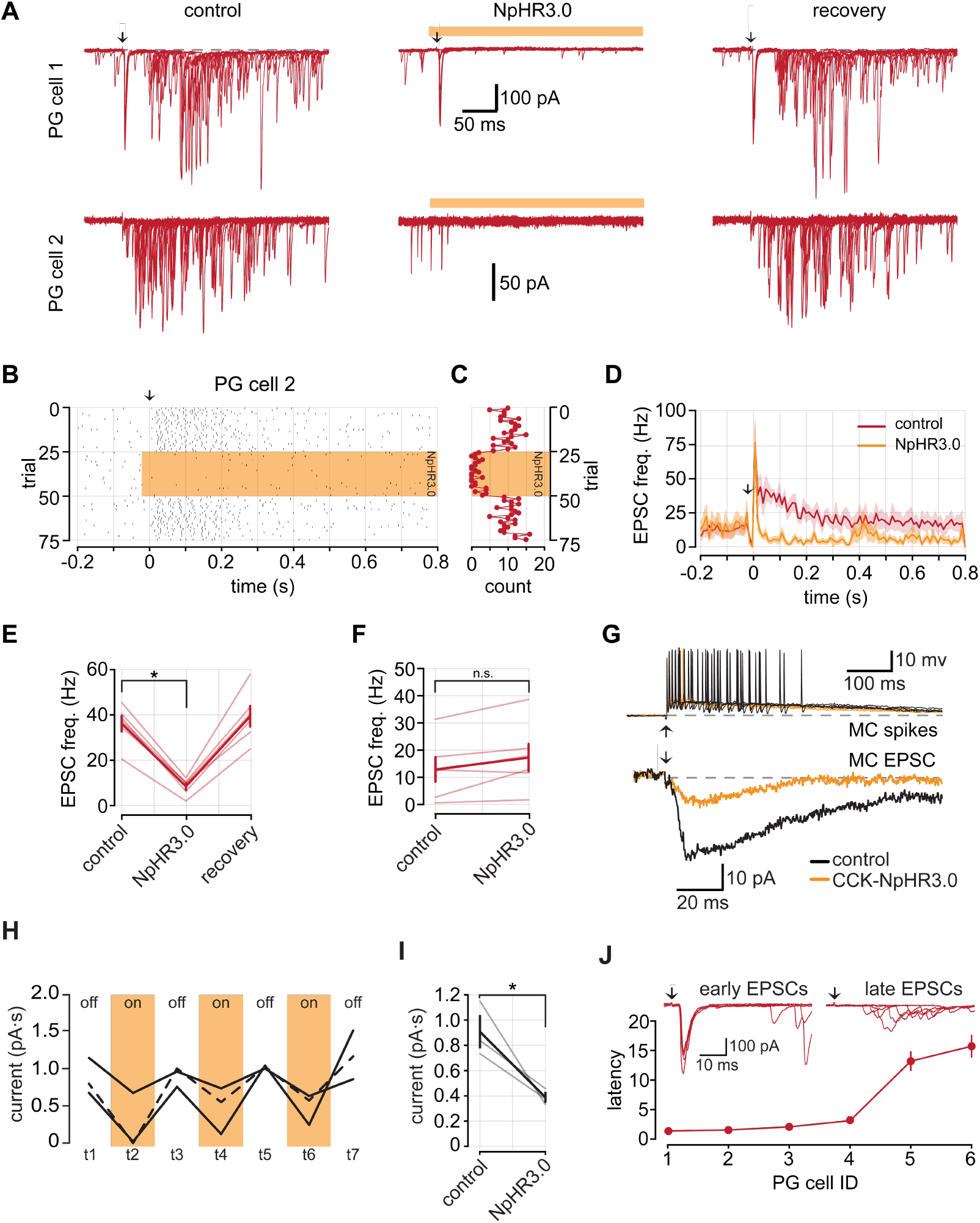
Evoked EPSCs in type II PG cells require activation of CCK-expressing TCs. **A**. Example EPSCs evoked by OSN stimulation in two PG cells. Each panel contains five overlaid trials. Downward arrows correspond to OSN stimulation. **B**. Raster plot of EPSCs from PG cell 2 in part *A*. The yellow bar indicates the presence of 590 nm light to suppress CCK-expressing cells with NpHR3.0. **C**. EPSC count of each trial in *B* in the 300 ms immediately following OSN stimulation. **D**. Summary PSTH of EPSC frequency across all PG cells (*n* = 6). The red trace is control trials; the yellow trace is trials with the LED on. Note: the drop in EPSC frequency right at OSN stimulation reflects obscuring of the EPSCs by the stimulus artifact; the jump in EPSC frequency that follows reflects direct OSN input to type II PG cells. **E**. Summary data of evoked EPSC frequency in the 300 ms following ON stimulation. **F**. Summary of baseline EPSC frequency during control and NpHR3.0 trials (12.93 ± 4.57 Hz control trials vs. 17.31 ± 5.08 Hz NpHR3.0 trials; *P* = 0.094; *Wilcoxon signed rank test*). **G**. *Top*, evoked spikes recorded in an MC under control conditions and during TC inactivation. *Bottom*, example excitatory currents recorded from the same MC. TC inactivation reduced the size of the current. **H**. Excitatory current area measurements from three MCs on interleaved trials of TC inactivation with NpHR3.0. The dashed line represents the MC in part *G*. **I**. Summary data of effects on TC inactivation on OSN-evoked excitatory currents measured in MCs. **J**. *Top*, example EPSCs from two different PG cells, one that received early as well as late input (*left*) and second that only received late excitatory input (*right*). *Bottom*, plot of the latency to the onset of the first EPSC across six PG cells.

One potential caveat with our results showing that TC suppression reduces excitation of PG cells is the possibility that the effect was an indirect result of TC suppression on MC excitation. Because eTCs are involved in a multi-step OSN-to-eTC-to-MC pathway for activating MCs (De Saint Jan et al., 2009; Najac et al., 2011; Gire et al., 2012), suppressing activation of CCK-expressing TCs could reduce MC excitation, which in turn reduces MC-to-PG cell signaling (Najac et al., 2015). Indeed, we found in a limited number of recordings in MCs that TC inactivation retrial-dependent and reversible reduction in MC excitatory current (56.52 ± 16.23% reduction in integrated charge; *n* = 3, *p* = 0.021; *Student’s T-test*; **Figure 4G-I**), consistent with TC inactivation suppressing MC activation. Thus, the reduced frequency of evoked EPSCs in PG cells during optogenetic suppression of TCs could be explained either because TCs account for the majority of direct excitatory synaptic inputs onto PG cells and/or because TC activation is required for MC excitation (see Discussion).

In previous physiological studies, Type II PG cells were defined to be PG cells that, upon OSN stimulation, receive feedforward excitation from eTCs (Shao et al., 2009). Furthermore, in contrast to Type I PG cells, Type II PG cells were defined *not* to receive direct synaptic input from OSNs. Our recordings suggested that at least some type II PG cells receive direct OSN input that can precede eTC-driven input (**Figure 4A**; PG cell 1). In three of the six cells included in our dataset we observed short latency (< 2.5 ms) EPSCs consistent with monosynaptic OSN input (**Figure 4J)**. Moreover, in these cells, NpHR3.0-mediated inactivation of TCs did not affect the short-latency early EPSC (**Figure 4A,D**), further consistent with these events reflecting direct OSN input initial OSN-driven EPSC. Thus, our results suggest that Type II PG cells can be excited by both eTCs and OSN inputs.

### eTCs provide direct input to PG cells but not eTCs

Thus far, our optophysiological experiments indicate that CCK-expressing TCs mediate strong excitation of PG cells across a variety of conditions while providing little direct excitatory synaptic input onto eTCs. We next tested, using dual-cell recordings in rat OB slices, whether similar conclusions applied to the presynaptic function of eTCs, which are a subset of CCK-expressing cells. Here, we recorded from one eTC in current-clamp configuration and either another eTC or a PG cell in voltage-clamp mode. We delivered depolarizing current injections to the eTC in current-clamp configuration to generate trains of action potentials (**Figure 5A**). Occasionally, stimulation of an eTC was sufficient to evoke a glomerulus-wide long-lasting depolarization (LLD; Carlson et al., 2000); for our analysis we did not consider these trials because we could not be certain of the origin of any excitatory signal that we measured during an LLD. In dual-cell recordings in which both cells were eTCs, we found that depolarization of one eTC resulted in small-amplitude but prolonged inward currents (**Figure 5Ai**). These slow eTC-to-eTC excitatory currents may reflect an extrasynaptic signaling mechanism, as has been described for eTC-to-MC transmission (Gire et al., 2012, 2019). Importantly, we found no evidence that eTC stimulation evoked fast EPSCs in the eTC-eTC pairs that is consistent with typical synaptic connections (15.28 ± 2.46 Hz baseline vs. 13.79 ± 2.30 Hz following eTC stimulation; *n* = 25 pairs; *P* = 0.023; *Wilcoxon signed rank test;* **Figure 5A,C,D**). In contrast, direct stimulation of an eTC in eTC-PG cell pairs caused a large increase in the frequency of fast EPSCs in the PG cell (11.40 ± 3.61 Hz baseline vs. 24.09 ± 2.40 Hz eTC stimulation; *n* = 6 pairs; *P* = 0.031; *Wilcoxon signed rank test*; **Figure 5C,D**), as expected for direct eTC-to-PG cell contacts (Hayar et al., 2004a; Najac et al., 2015; Tavakoli et al., 2018). Notably, in our analysis of fast EPSCs in the eTC-eTC pairs, we found that direct stimulation of an eTC not only failed to induce fast EPSCs, but in fact, induced a small but significant reduction in the frequency of fast EPSCs (**Figure 5D**). The cause of the reduced EPSC frequency could be modulation of glutamate release from OSNs (e.g., via GABA_B_ receptors following activation of PG cells) (Nickell et al., 1994; Aroniadou-Anderjaska et al., 2000; Wachowiak et al., 2005). Alternatively, it could potentially be an artifact of the higher current noise associated with the evoked slow currents (**Figure 5Ai**), which impacted the EPSC event detection. We did not further investigate this effect.

**Figure 5.**
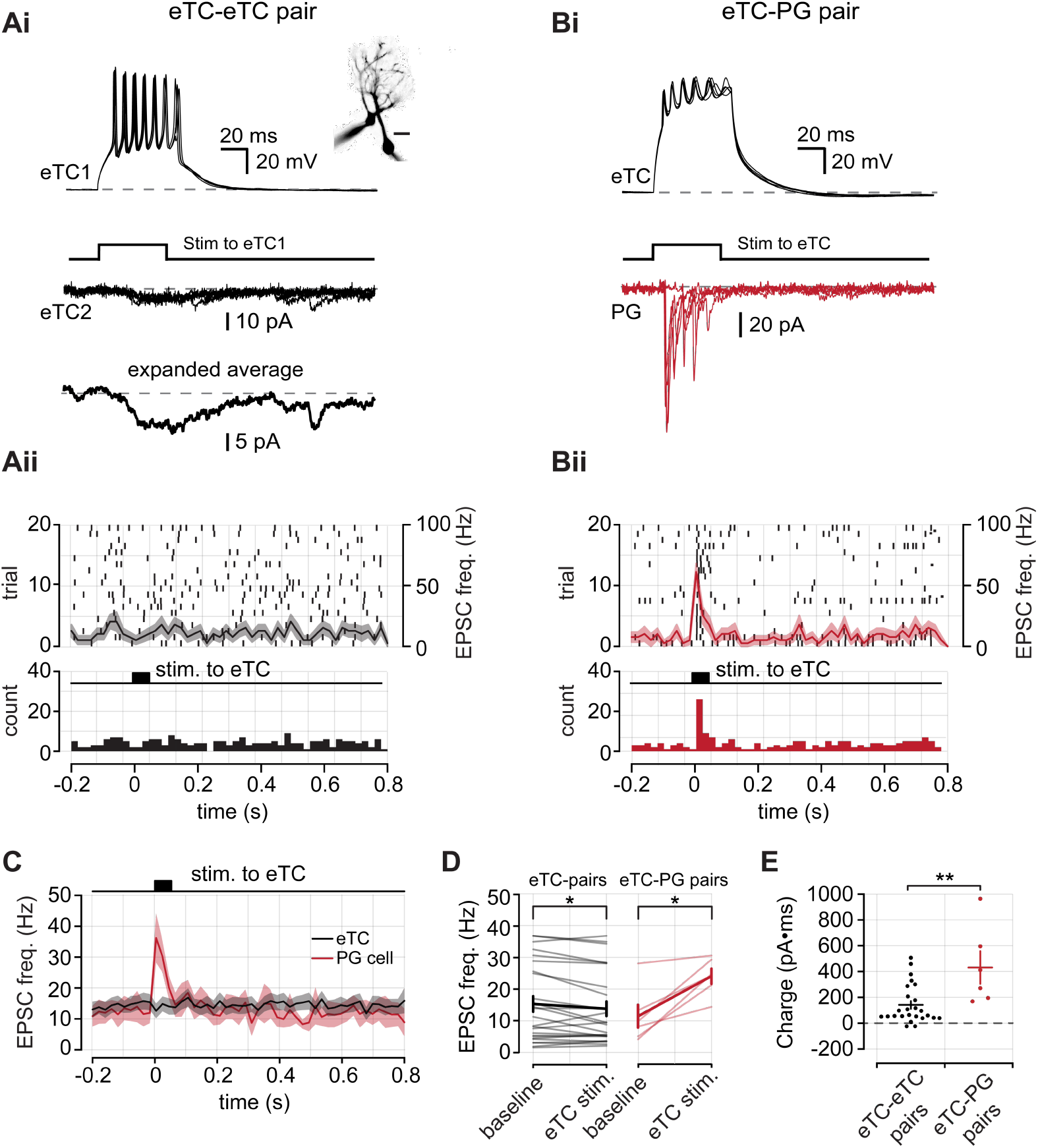
Pair-cell recordings show that eTCs provide direct input to PG cells, but not other eTCs. **A**. (**i**) Example eTC-eTC pair recording in rat olfactory bulb slices. eTC1 is recorded in current-clamp and depolarized to generate spikes. Excitatory currents are recorded from eTC2 in voltage-clamp configuration. Three consecutive sweeps are shown in both cells as well as (*bottom*) the average response in eTC2. Note that the evoked responses are dominated by a small-amplitude prolonged current. A cell-fill image of the eTC-eTC pair is shown in the inset. Scale bar = 20 μm. (**ii**) *Top*, raster plot of rapid EPSCs detected in eTC2 from part *i* across 20 trials. EPSCs were detected as in previous figures. The black curve is the mean PSTH and the shaded area is s.e.m. *Bottom*, pooled bin counts for 20 trials. **B**. Same as part *A*, but for an eTC-PG cell pair. The eTC was depolarized in current-clamp configuration to generate spikes and EPSCs were measured from the PG cell in voltage-clamp mode. **C**. Mean rapid EPSC PSTH from 25 eTC-eTC pairs and 6 eTC-PG cell pairs. **D**. Summary data of the mean EPSC frequency in the baseline period and the 80 ms following eTC stimulation. **E**. Summary data comparing the integrated charge in postsynaptic eTCs and PG cells (141.65 ± 30.04 pA•ms in eTC-eTC pairs vs. 432.49 ± 126.60 pA•ms in eTC-PG cell pairs; *n* = 25 eTC-eTC pairs and 6 eTC-PG cell pairs; *P* = 0.010; *Wilcoxon rank sum test*).

The pair-cell recordings provided excellent evidence that eTCs fail to synaptically excite other eTCs, although they can drive a prolonged excitatory current in eTCs. To determine the potential impact of the slow eTC-to-eTC currents, we compared the integrated charge associated with the slow currents with that associated with the more typical barrages of evoked EPSCs in PG cells (**Figure 5Bi**). To capture the entirety of the slow EPSC, we used a 100 ms integration window that was aligned to the onset of the depolarizing current injection in the presynaptic cell. For both cell types, charge measurements were obtained from composite traces that contained at least 15 trials. We found that for a similar number of presynaptic spikes (4.56 ± 0.33 spikes/trial in eTC-eTC pairs and 4.43 ± 0.52 spikes/trial in eTC-PG cell pairs; *P* = 0.940; *Wilcoxon rank sum test*) excitation between eTCs is weaker than between eTCs and PG cells (141.65 ± 30.04 pA•ms in eTC-eTC pairs, *n* = 25, vs. 432.49 ± 126.60 pA•ms in eTC-PG cell pairs, *n* = 6; *P* = 0.010; *Wilcoxon rank sum test***; Figure 5E**). We did however observe a few eTC pairs where the integrated charge was similar to that measured in eTC-PG cell pairs. These instances indicate that under certain conditions, perhaps related to spatial proximity or glutamate clearance properties, eTCs might signal to other eTCs with increased efficacy.

### Optical manipulation of intraglomerular inhibition at eTCs

Our studies in CCK-NpHR3.0 mice, together with the pair-cell recordings, provided information about the role of TCs in driving excitation of different cell-types in the glomerular layer including PG cells. In the last part of this study, we considered the reverse step – the inhibition of TCs by PG cells. The glomerular layer includes PG cells that differ in expression of GAD isoforms, either GAD65 or GAD67, and also in the number of glomeruli to which they send apical dendrites (Parrish-Aungst et al., 2007; Kiyokage et al., 2010). eTCs can also receive GABAergic input from short-axon cells located in the glomerular layer (Liu et al., 2013; Banerjee et al., 2015). We wondered what contribution GAD65-expressing PG cells have on inhibition of eTCs. GAD65-expressing PG cells are the most numerous class of PG cells and generally have dendritic arbors that are confined to one glomerulus (Parrish-Aungst et al., 2007; Kiyokage et al., 2010). Hence they mediate a unique form of local inhibition.

We generated GAD65-tdTomato mice, in which we confirmed somatic reporter expression interspersed throughout the glomerular layer (**Figure 6A**), which was consistent with previous reports of GAD65 expression (Parrish-Aungst et al., 2007; Kiyokage et al., 2010; Whitesell et al., 2013; Sun et al., 2020). We also observed dense reporter expression in the EPL that likely arose from the apical dendrites of granule cells that also express GAD65; however, granule cells do not contact eTCs, which lack lateral dendrites. We then generated GAD65-NpHR3.0 mice to selectively manipulate GAD65-expressing cells. NpHR3.0 activation resulted in strong membrane hyperpolarizations in PG cells (**Figure 6B**) and completely blocked spikes driven by OSN stimulation (**Figure 6C-D**). We next measured the effect of inactivating GAD65-expressing cells while recording OSN-evoked inhibitory postsynaptic currents (IPSCs) in eTCs (*V*_*hold*_ = ≥ 0 mV; **Figure 6E**). For the current recordings, blocks containing 10 trials with or without light pulses were alternated. When slices were illuminated, the evoked IPSCs in eTCs were reduced by about half (43.48 ± 4.76% reduction in integrated charge; *n* = 5; *P* < 0.001; *Paired T-test;* **Figure 6F**). However, some IPSC remained and might reflect either incomplete suppression of activity in GAD65-expressing PG cells or evoked inhibitory input from other GABAergic cell-types in the glomerular layer. In four of five recordings, the initial IPSC amplitude was recovered during the second block of control trials (88.76 ± 7.71% of initial control, *n* = 4), indicating that the reduced IPSC during the light pulses was not a result of run-down.

**Figure 6.**
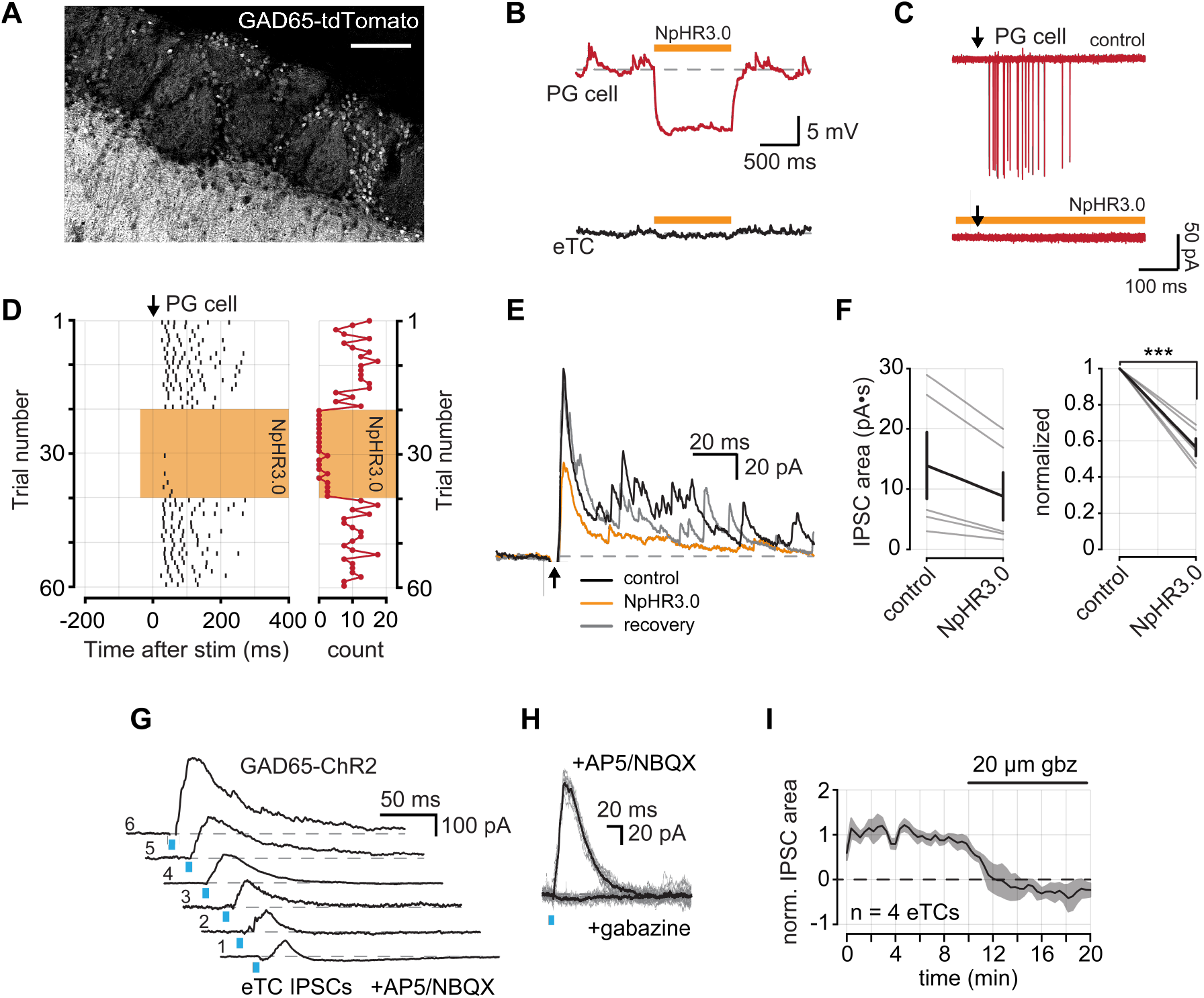
GAD65-expressing PG cells inhibit eTCs. **A**. GAD65 expression in a tdTomato reporter mouse line. GAD65 is expressed in the soma of PG cells that surround glomeruli. GAD65 expression is also observed in the EPL, likely reflecting granule cell apical dendrites. Scale bar = 100 μm. **B**. Example membrane voltage responses from a PG cell (top) and an eTC (bottom) in a GAD65-NpHR3.0 mouse in response to a 1 s pulse of 590 nm light. **C**. Example OSN-evoked (8 µA stimulus) spikes from a PG recorded in LCA configuration. Five overlaid trails under control conditions and during NpHR3.0 activation. Downward arrow corresponds to OSN stimulation. **D**. *Left*, raster plot of evoked spikes per trial from the cell in part C. *Right*, a plot of the number of spikes following each OSN stimulation. **E**. Example IPSCs evoked by OSN stimulation recorded from an eTC (V_*hold*_ = +35 mV) under control conditions, during NpHR3.0 activation, and following recovery. Each trace is an average of 10 consecutive trials in each condition. **F**. *Left*, summary of OSN evoked IPSC areas. *Right*, summary of normalized IPSC areas (43.48 ± 4.76% reduction; *n* = 5; *P* < 0.001; *Student’s T-test*). **G**. Example IPSCs recorded from six eTCs following 470 nm LED light stimulation in OB slices prepared from GAD65-ChR2 mice. **H**. IPSCs were eliminated by bath application of gabazine. Light-colored traces are individual trials and dark is the mean of all trials. **I**. Time course of IPSC block with gabazine from four eTCs. IPSC areas are normalized to the mean control area for each cell.

As a further confirmation that GAD65-expressing PG cells provide inhibitory input to eTCs, we also measured evoked currents in eTCs from bulb slices prepared from GAD65-Channelrhodopsin2 (ChR2) mice. Recordings were conducted in the presence of the glutamate receptor antagonists NBQX (20 µM) and DL-APV (50 µM) to eliminate the possibility that simultaneously depolarizing a large population of GAD65-expressing cells may have off-target network effects. In each test eTC (*n* = 6), brief light pulses (5 ms at 423 nm) evoked large IPSCs (**Figure 6G**). We confirmed that these currents were GABA_A_ receptor-mediated by their sensitivity to the antagonist gabazine (20 µM; **Figure 6H,I**; *n* = 4). Taken together, our findings in GAD65-NpHR3.0 and GAD65-ChR2 mice provide novel optogenetic evidence that local GAD65-expressing PG cells strongly inhibit eTCs.

## Discussion

Prior studies have established that a subset of TCs, the eTCs, play a central role in mediating both inhibition and excitation within OB glomeruli. The inhibition is driven by reciprocal dendrodendritic contacts between eTCs and GABAergic PG cells (Hayar et al., 2004a; Hayar et al., 2005; Murphy et al., 2005), while excitation arises from a feedforward pathway wherein OSNs first excite eTCs which, in turn, excite MCs (OSN-to-eTC-to-MC; De Saint Jan et al., 2009; Najac et al., 2011; Gire et al., 2012). In this study, we used optophysiological tools and pair-cell recordings to delineate a number of novel aspects of the connectivity between TCs and PG that are important for understanding sensory processing in glomerular networks.

### Neural connectivity within olfactory bulb glomeruli

Our first new finding about glomerular connections was that eTCs specifically, and TCs more generally, <u>fail</u> to make chemical synaptic connections onto eTCs. In recordings from eTCs in CCK-NpHR3.0 mice, we found that optogenetically suppressing TCs had no effect on the frequency or amplitude of fast EPSCs that are mediated by typical AMPA receptor glutamatergic connections, both under spontaneous conditions and also following stimulation of OSN axons. If TC-to-eTC connections were prevalent, suppression of TCs should have reduced the EPSCs in some capacity. The absence of an effect was not because our optogenetic methods were ineffective in suppressing TCs, since our control experiments indicated strong light-evoked suppression of TC output (**Figure 1**). The lack of effect was also not because TCs were not active during the conditions of these experiments, since TC suppression was highly effective in reducing EPSCs in PG cells reflecting direct eTC-to-PG cell contacts (Hayar et al., 2004a). Further arguing that eTCs do not make strong connections onto eTCs was that, in pair-cell recordings, eTC stimulation did not drive rapid EPSCs in eTCs, even though such stimulation was effective in doing so in eTC-PG cell pairs. Interestingly, we did find in the eTC-eTC pairs that eTC stimulation resulted in a low-amplitude slow excitatory current that lasted ∼100 ms, suggesting that TCs may at a minimum weakly excite eTCs through other modes of signaling. The slow eTC currents resembled slow currents that have been reported in eTC-to-MC pairs that result from the spill-over of glutamate at eTC-to-PG cell synapses (Gire et al., 2019), suggesting the possibility of a similar underlying mechanism.

The question of whether there are glutamatergic synaptic connections between TCs/eTCs has been addressed previously using ultrastructural methods. Kosaka and Kosaka (2005) reported the presence of dendrodendritic synaptic connections between excitatory dendrites within glomeruli of mouse OB, potential TC-to-TC contacts, although they did not quantify their number nor determine cell types. Prior studies from our lab (Bourne and Schoppa, 2017) also provided evidence for such connections in glomeruli of rat OB, although their frequency of occurrence was very low. The evidence from our physiological studies here for negligible TC-to-eTC connections fits well with the latter, more quantitative ultrastructural results, and it is gratifying that the two analysis methods, which rely on different assumptions, led to similar conclusions. An additional point that should be made about the physiological analysis is that it, unlike the ultrastructural work, was agnostic about the location and specific type of glutamatergic connections between eTCs being tested. There is now clear evidence that at least some subtypes of TCs can form local axon collaterals within OB (Ojima et al., 1984; Orona et al., 1984; Igarashi et al., 2012; Sun et al., 2020), which in principle could form axodendritic contacts onto eTCs either within or outside of glomeruli. Our results showing that TCs do not mediate fast EPSCs in eTCs argues against both axodendritic and dendrodendritic contacts between TCs/eTCs.

The second new result from our study regards the previously-described dendrodendritic connections that eTCs make onto Type II PG cells (Hayar et al., 2004a). While it has been unambiguous that these connections exist, it has been uncertain how much these inputs contribute to excitation of PG cells versus other sources such as MCs (Najac et al., 2015) and centrifugal inputs from olfactory cortical regions (Boyd et al., 2012; Markopoulos et al., 2012; Brunert et al., 2016). The fact that optogenetic suppression of TCs resulted in 70-75% decrease in the spontaneous and evoked EPSCs in PG cells is consistent with the majority of excitatory inputs into Type II PG cells reflecting inputs from TCs under these conditions. There are some limitations with these results. For example, when considering MCs as a potential source of inputs onto Type II PG cells, at least part of the reduction in evoked EPSCs in PG cells following suppression of TCs could in principle have been due to reduced feedforward excitation of MCs that contact PG cells rather than reduced TC input. Nevertheless, we believe that our results do enable us to make a strong conclusion about the role of TCs in driving excitation of PG cells. In both scenarios – whether the reduced EPSCs due to NpHR3.0 activation was a direct result of suppressing TC-to-PG inputs or resulted indirectly from suppressing feedforward excitation of MCs – TC activation was a necessary step in driving most of the excitation of PG cells. Our results explicitly exclude a pathway involving direct inputs from OSNs onto MCs (Najac et al., 2011) as a dominant mechanism of exciting PG cells (i.e., an OSN-to-MC-to-PG cell pathway).

A third and final new mechanism pointed to by our optophysiological studies regards the inhibitory synaptic input that eTCs receive from GABAergic interneurons. Prior studies (Parrish-Aungst et al., 2007; Kiyokage et al., 2010) have established that the glomerular layer includes GABAergic PG cells that can be biochemically segregated by expression of different isoforms of GAD (65 or 67). These biochemically-defined PG cells also differ in their anatomy. Most GAD65-expressing PG cells are “uniglomerular”, having an apical dendrite confined to one glomerulus, while most GAD67-expressing PG cells have apical dendrites that extend into multiple glomeruli (Kiyokage et al., 2010). Many GAD67-expessing cells also have axons that can extend long distances across the glomerular layer. Based on the fact that suppressing GAD65-positive cells in GAD65-NpHR3.0 mice resulted in a ∼50% reduction in evoked IPSCs in eTCs, we conclude that <u>at least</u> that much of the inhibition under this condition was provided by GAD65-positive, uniglomerular PG cells. A broader conclusion can also be made If we consider the type of PG cells that we were studying in the analysis of TC-to-PG cell connectivity (see above). Because the post-synaptic PG cells in those studies were uniglomerular, our results taken together argue that there are strong reciprocal connections between eTCs and GAD65-positive, uniglomerular PG cells.

### Functional implications

What are the functional implications of our new findings about the microcircuitry of glomeruli in OB? If we assume a model in which the output of a glomerulus is determined by the balance between excitation and inhibition, our combined results point toward a scenario in which odors should generally fail to produce a glomerular output due to an unfavorable E/I balance. We failed to observe synaptic connections amongst TCs/eTCs, which could underlie recurrent excitation, while, at the same time, eTCs make strong reciprocal connections with a specific class of inhibitory PG cells. Our conclusion, that the glomerular E/I balance should generally be weighted toward inhibition, is also supported by the available literature characterizing odor-evoked responses in MCs. For example, MCs are much more narrowly tuned to odors than are OSNs (Tan et al., 2010; Kikuta et al., 2013), which at least in part reflects odor-evoked inhibition (Yokoi et al., 1995). It should be pointed out that, in tying our results about the E/I balance at the level of TCs/eTCs to the properties of output MCs, there is an assumption that the E/I balance in TCs/eTCs should impact MC activation. This is well-supported by the available literature. Brain slice studies have shown that the majority of the excitatory current in MCs following OSN stimulation is driven by eTCs (Gire et al., 2012; Vaaga and Westbrook, 2016). Thus, the activation status of eTCs, which is dictated by their E/I balance, should be a major contributor to MC activation.

If the E/I balance in a glomerulus generally favors inhibition, might there be conditions in which the E/I balance can become more favorable? Some clue to this was provided by recent studies that analyzed eTC-to-MC feedforward excitation across a range of stimulus conditions (Gire et al., 2019). These studies found that weak stimuli (few spikes in eTCs) produced very little eTC-to-MC excitation, but this excitation rose in a highly supralinear fashion with stimulus strength. Such changes, which may in part reflect the dynamics of extrasynaptic glutamate that underlies feedforward excitation, could be critical for allowing MCs to be excited by a select set of “strongest” stimuli, contributing to the narrow tuning of MCs (see above). We suggest that similar mechanisms could be revealed if we were to further analyze the slow, non-synaptic excitatory signals that were observed in eTCs upon eTC stimulation (see above). While these currents were small under the conditions tested here, a greater number of eTC spikes and/or the concerted activity of many eTCs could amplify these signals. Clearly, more studies are needed to understand eTC-to-eTC signaling under a range of stimulus conditions, as well as how they are balanced with inhibition.

## Methods

### Animals

CCK-Cre mice (Chhatwal et al., 2007) (Jackson Laboratory stock #011086) were maintained as heterozygous breeding pairs. Heterozygous CCK-Cre offspring were then crossed with LSL-eNpHR3.0-EYFP mice (Madisen et al., 2012) (Jackson Laboratory stock #014539). The resulting male and female offspring were used for optogenetic experiments. CCK-Cre mice were also crossed with LSL-tdTomato mice (Madisen et al., 2010) (Jackson Laboratory stock #07909) to generate reporter animals to verify CCK expression. GAD65-Cre mice (Taniguchi et al., 2011) (Jackson Laboratory stock #019022) were also maintained as heterozygous breeding pairs and crossed with LSL-eNpHR3.0-EYFP or LSL-ChR2-EYFP mice (Madisen et al., 2012). GAD65-Cre mice were also crossed with LSL-tdTomato mice to make reporter animals. Male and female Sprague Dawley rats at P8 to P20 (Charles River Laboratories) were used for non-optogenetic experiments. Animals were housed on a 12 hr light-dark schedule and fed ad libitum. All experiments were conducted under protocols approved by the Animal Care and Use Committee of the University of Colorado, Anschutz Medical Campus.

### Slice preparation

Acute horizontal OB slices (300–330 μm) were prepared following isoflurane anesthesia and decapitation. For experiments using genetic expression of light-gated ion channels, adult mice (> 6 weeks) were used to allow for sufficient channel expression. Both OBs were rapidly removed and placed in oxygenated (95% O_2_, 5% CO_2_) ice-cold solution containing the following (in mM): 72 sucrose, 83 NaCl, 26 NaHCO_3_, 10 glucose, 1.25 NaH_2_PO_2_, 3.5 KCl, 3 MgCl_2_, and 0.5 CaCl_2_ adjusted to 295 mOsm. OBs were separated into hemispheres with a razor blade and attached to a stage using adhesive glue (Loctite 404) applied to the ventral surface of the tissue. Slices were cut using a vibrating microtome (Leica VT1000S) and were incubated in a holding chamber for 30 min at 32°C. Subsequently, the slices were stored at room temperature.

### Electrophysiology

All electrophysiology experiments were conducted under an upright Zeiss Axioskop2 FS Plus microscope (Carl Zeiss MicroImaging) fitted with differential interference contrast (DIC) optics, video microscopy under control of Slidebook software (3i), and a CCD camera (Hamamatsu). Cells were visualized and identified with 10x or 40x Zeiss water-immersion objectives. All recordings were performed at 32– 35°C.

The base extracellular recording solution contained the following in mM: 125 NaCl, 25 NaHCO_3_, 1.25 NaHP0_4_O, 25 glucose, 3 KCl, 1 MgCl_2_l, and 2 CaCl_2_ (pH 7.3 and adjusted to 295 mOsm), and was oxygenated (95% O_2_, 5% CO_2_). The base intracellular pipette solution for whole-cell current-clamp recordings contained the following in mM: 125 K-gluconate, 2 MgCl_2_, 0.025 CaCl_2_, 1 EGTA, 2 Na_3_ATP, 0.5 Na_3_GTP, and 10 HEPES (pH 7.3 with KOH, osmolarity adjusted to 215 mOsm) (Zak et al., 2015). For some current recordings, K-gluconate in the pipette solution was replaced with an equimolar amount of cesium methanosulfonate, as well as the sodium channel blocker QX-314 (10 mM) to block action potentials. All For whole-cell current-clamp recordings from eTCs, 30 mM glutamic acid was added to the pipette to prevent the run-down of evoked neurotransmitter release (Ma and Lowe, 2007). Loose cell-attached (LCA) recordings from eTCs were made with a pipette that contained 150 mM NaCl. All whole-cell recordings included 100 μM Alexa Fluor 488 or Alexa Fluor 594 (Invitrogen) in the pipette solution to allow for visualization of cell processes. Fluorescence measurements were performed under whole-field epi-illumination using a DG-4 light source (Sutter Instruments). Signals were detected by a Cool-Snap II HQ CCD camera (Photometrics) under control of Slidebook software.

Borosilicate glass patch pipettes (World Precision Instruments) were pulled to a resistance of 4–6 MΩ for eTCs, 6–8 MΩ for PG cells, and 3– 4 MΩ for MCs using an upright puller (Narishige). Current and voltage signals in the single- and pair-cell experiments were recorded with a Multiclamp 700B amplifier (Molecular Devices), low-pass filtered at 1.8 kHz using an eight-pole Bessel filter, and digitized at 10 kHz using a Digidata 1322A (Axon Instruments) digital interface. Data were acquired using AxographX software on an Apple MacPro computer.

Cell identity was determined in part by visualizing Alexa Fluor 488-mediated or Alexa Fluor 594-mediated fluorescence signals. eTCs were distinguished from PG cells by their position in the inner half of the glomerular layer, their relatively large, spindle-shaped somata (∼10 μm in diameter), a single highly branched apical dendrite, lack of lateral dendrites, and relatively low input resistance (∼0.2 GΩ; Hayar et al., 2004b). PG cells were identified by their small soma (<10 μm in diameter) and high input resistance (>0.8 GΩ; Hayar et al., 2004b; Murphy et al., 2005; Shao et al., 2009). Test PG cells were all uniglomerilar, with an apical dendrite that extended to just one glomerulus (Kiyokage et al., 2010). We only considered PG cells that displayed bursts of EPSCs either spontaneously or in response to OSN stimulation, consistent with Type II cells (Kosaka et al., 1998; Shao et al., 2009). MCs were easily identified by their position in the mitral cell layer and distinctive dendritic arborizations.

Stimulation of OSN axons was performed using a broken-tip patch pipette (5–10 μm in diameter) placed in the olfactory nerve layer, ∼50– 100 μm superficial to the glomerular layer. Current injections were delivered by a stimulus isolator (World Precision Instruments) under control of a TTL output from AxographX software. Weak intensities of electrical stimulation were used (5– 50 μA). eTCs or PG cells that were chosen were associated with glomeruli at the surface of the slice. Stimulus artifacts in many of the illustrated traces have been blanked or truncated. Optogenetic stimulation of NpHR3.0 was performed using a 140 mW 590 nm collimated LED (ThorLabs) with epi-field illumination through a 40x objective under control of a TTL output from AxographX software. ChR2 stimulation was performed using a 760 mW 470 nm LED (ThorLabs). For displayed voltage-clamp traces of eTCs where NpHR3.0 is activated (**Figure 2D**), current artifacts were removed by subtracting a median filtered signal from each trial individually.

### Imaging

Confocal images of CCK-tdTomato and GAD65-tdTomato expression were acquired by scanning laser confocal microscope (Olympus Fluoview FV1000). Maximum projection images were generated from z-stacks at 1 µm intervals in a single field of view using Fiji (Schindelin et al., 2012).

### Experimental design and statistical methods

Evoked and spontaneous EPSCs were detected using a variable amplitude template with a 0.5 ms rise and 2 ms decay, consistent with typical AMPA receptor kinetics (Schoppa, 2006; Tyler et al., 2007; Grubb et al., 2008). EPSCs with amplitudes < 2.5 standard deviations of baseline noise were rejected. Data were analyzed using AxographX and custom MATLAB scripts (Mathworks). Data throughout are expressed as mean ± s.e.m.

Peristimulus time histograms (PSTH) of EPSC frequency and spike rates were computed by calculating the mean frequency in 20-50 ms time bins from ≥ 10 sweeps. Significance was determined using two-tailed non-parametric tests, a *Wilcoxon signed rank test* for matched pairs, or a *Wilcoxon rank sum* test for non-matched samples. Paired *T-*tests were used in some matched pair comparisons due to a smaller sample size when *n* < 6. Throughout, a value of *P* < 0.05 was considered significant.

## Acknowledgements

This work was supported by NIH grant R01 DC006640 to NES and NIH fellowship F31 DC015938 to JDZ.

## Author Contributions

JDZ and NES designed the experiments. JDZ acquired and analyzed the data. JDZ and NES wrote the manuscript.

